# The transcriptional aftermath in two independently formed hybrids of the opportunistic pathogen *Candida orthopsilosis*

**DOI:** 10.1101/2020.03.27.012922

**Authors:** Hrant Hovhannisyan, Ester Saus, Ewa Ksiezopolska, Toni Gabaldón

## Abstract

Interspecific hybridization can drive evolutionary adaptation to novel environments. The *Saccharomycotina* clade of budding yeasts includes many hybrid lineages, and hybridization has been proposed as a source for new pathogenic species. *Candida orthopsilosis* is an emerging opportunistic pathogen for which most clinical isolates are hybrids, each derived from one of at least four independent crosses between the same two parental lineages. To gain insight on the transcriptomic aftermath of hybridization in these pathogens, we analyzed allele-specific gene expression in two independently formed hybrid strains, and in a homozygous strain representative of one parental lineage. Our results show that the effect of hybridization on overall gene expression is rather limited, affecting ~4% of the studied genes. However, we identified a larger effect in terms of imbalanced allelic expression, affecting ~9.5% of the heterozygous genes in the hybrids. Some of these altered genes have functions related to pathogenicity, including zinc transport and superoxide dismutase activities. Additionally, the number of shared genes with imbalanced expression in the two independently formed hybrids was higher than random expectation, suggesting selective retention. While it remains unclear whether the observed imbalanced genes play a role in virulence, our results suggest that differences in allele-specific expression may add an additional layer of phenotypic plasticity to traits related to virulence in *C. orthopsilosis* hybrids.

**Importance:** How new pathogens emerge is an important question that remains largely unanswered. Some emerging yeast pathogens are hybrids originated through the crossing of two different species, but how hybridization contributes to a higher virulence is unclear. Here we show that hybrids selectively retain gene regulation plasticity inherited from the two parents, and that this plasticity affects genes involved in virulence.

## Introduction

The incidence of human fungal infections has steadily increased during the past decade, leading to recognition of their relevance in global epidemiology (Pfaller and Diekema 2007). Numerous factors may underlie this growing prevalence including, among others, increased number of immunocompromised individuals (elderly people, neonates, HIV, patients, etc) (Pfaller and Diekema 2007), emergence of drug resistance associated with extensive use of antimycotic agents (Pfaller et al. 2009; Ksiezopolska and Gabaldón 2018), globalization (Mixão and Gabaldón 2018; Callaghan and Guest 2015), and climate change (Garcia-Solache and Casadevall 2010). The rising incidence of mycoses is also coupled to the identification of a larger number of etiological agents, including so called emergent pathogens that are increasingly identified in the clinics (Gabaldón, Naranjo-Ortíz, and Marcet-Houben 2016; Papon et al. 2013). In this context, hybridization between different species or lineages has been identified at the origin of several emerging yeast pathogens (Mixão et al. 2019; Schröder et al. 2016; Pryszcz et al. 2015, 2014; Wu et al. 2015; Mixão and Gabaldón 2018).

For some fungal hybrid pathogens the two parental species have been identified, as in the case of *Cryptococcus neoformans* × *Cryptococcus deneoformans* (Boekhout et al. 2001), and *Cryptococcus neoformans* × *Cryptococcus gattii* (Bovers et al. 2008). For others, one or both (or possibly numerous) parentals are still unknown. For example, for *Candida orthopsilosis* only one of the two putative parental lineages has been identified (Pryszcz et al. 2014; Schröder et al. 2016), which constitutes a minority (~7%) of the analyzed clinical strains. In the case of *Candida metapsilosis* or *Candida inconspicua* both parentals remain unknown (Pryszcz et al. 2015; Mixão et al. 2019), as all analyzed strains are hybrids. The fact that some parental species of hybrid fungal pathogens are never or rarely identified among clinical isolates suggests that some human pathogens have arisen from non-pathogenic parental organisms (Pryszcz et al. 2014; Mixão and Gabaldón 2018), or that the parental lineages have been out-competed by their more adapted pathogenic hybrid descendants (J. R. Depotter et al. 2016; Jonge et al. 2013; J. R. L. Depotter et al. 2016). In either case, whether interactions between the two distinct sub-genomes contribute to emerging properties in the hybrid, such as increased ability to infect humans, is still poorly understood.

At the molecular level, hybridization results in a state often referred as a “genomic shock” (McClintock 1984), in which two diverged genomes which have evolved independently for a certain time are now sharing the same cellular environment. This coexistence can lead to alterations at several levels, including the genome (Dutta et al. 2017), the transcriptome (Cox et al. 2014), or the proteome (Hu et al. 2017), among others (Greaves et al. 2015). Advances in next-generation sequencing have facilitated the study of the genomic aftermath of hybridization, which involves phenomena such as large-scale genome duplications or deletions, homeologous recombination, and gene conversion leading to loss of heterozygosity (Mixão and Gabaldón 2018; Smukowski Heil et al. 2017; Dutta et al. 2017; Pryszcz et al. 2015; Marcet-Houben and Gabaldón 2015).

However, our understanding of the impact of hybridization at the transcriptomic level remains poorly characterized, with few studies performed on industrial or plant saprophyte hybrids (Metzger, Wittkopp, and Coolon 2017; Tirosh et al. 2009; Cox et al. 2014; X. C. Li and Fay 2017; Hovhannisyan et al. 2020). To date no study of the transcriptomic aftermath of hybridization has been performed in hybrid human pathogens, limiting our insights on how hybridization leads to emergent traits, including virulence. To fill in this gap, we here undertook a transcriptomic analysis of two independently formed hybrid strains from the emerging yeast pathogen *Candida orthopsilosis* (Pryszcz et al. 2014; Schröder et al. 2016). This yeast belongs to the CTG clade and is phylogenetically placed within the *C. parapsilosis sensu lato* species complex alongside with the other opportunistic pathogens *C. parapsilosis* (Pammi et al. 2013; Tóth et al. 2019) and *C. metapsilosis* (Tavanti et al. 2005). It has been shown that most (~93%) of the clinical isolates of *C. orthopsilosis* are hybrids between parentals with ~5.1% nucleotide divergence. As mentioned above, only one of the two parental lineages has been found among clinical isolates (Schröder et al. 2016). To date, the other partner in the hybridization remains unidentified. Notably, these two parental lineages have hybridized several independent times, giving rise to at least four distinct hybrid clades that differ in their levels and patterns of loss of heterozygosis (LOH, Schröder et al. 2016). Clade 1 comprises strains which undergone extensive LOH, and are thought to derive from a relatively ancient hybridization, while strains in clade 4 are the most heterozygous, with fewer LOH events and are thus assumed to result from a more recent event. Thus, *C. orthopsilosis* represents an appropriate model to study how hybridization impacts transcription in a natural hybrid pathogen, and whether parallel hybridization events result in similar transcriptomic interactions between the two parental sub-genomes.

To shed light on these questions, we conducted allele-specific expression (ASE) analysis using RNA-Seq of two hybrid strains of *C. orthopsilosis* each resulting from an independent hybridization event, that represent the two extremes of LOH extent, namely MCO456 (clade 1) and CP124 (clade 4). For comparison, we investigated the transcriptome of a highly homozygous strain belonging to one of the putative parental lineages (strain 90-125). To our knowledge this is the first description of the transcriptomic profiles between parental species in a hybrid yeast that is an opportunistic pathogen of humans.

## Materials and Methods

### Strains

We analyzed three diploid strains of *C. orthopsilosis* - MCO456, CP124, and 90-125, with the latter belonging to the lineage of one of the putative parentals of the two former hybrid species (Tavanti et al. 2005; Riccombeni et al. 2012; Pryszcz et al. 2014).

### Experimental conditions and RNA extraction

RNA extraction was performed on the samples growing at the exponential phase in rich yeast extract peptone dextrose medium (YPD) at 30°C.

Experiments were performed as follows:

First, we measured growth curves for each individual strain to delimit the mid-exponential growth phase. For this, each strain was plated on a YPD agar plate and grown form 3 days at 30°C. Single colonies were cultivated in 15 mL YPD medium in an orbital shaker (30°C, 200 rpm, overnight). Then, each sample was diluted to an optical density at 600nm (OD) of 0.2 in 50 mL of YPD and then grown for 3h at the same conditions. Then samples were diluted again to OD of 0.1 in 50 mL of YPD to start all experiments with a similar amount of cells. Cultures were grown at 30°C for 24 hours and OD was every 60 min with a TECAN Infinite M200 microplate reader. Upon reaching the mid-exponential phase, the protocol was repeated until all samples were growing at the exponential phase. Then cultures were centrifuged at 16 000g to harvest 3×10^8^ cells per sample. Total RNA from all samples was extracted using the RiboPure RNA Yeast Purification Kit (ThermoFisher Scientific) according to manufacturer’s instructions. Total RNA integrity and quantity of the samples were assessed using the Agilent 2100 Bioanalyzer with the RNA 6000 Nano LabChip Kit (Agilent) and NanoDrop 1000 Spectrophotometer (Thermo Scientific).

### RNA-Seq library preparation and sequencing

Sequencing libraries were prepared using the TruSeq Stranded mRNA Sample Prep Kit v2 (ref. RS-122-2101/2, Illumina) according to the manufacturer’s instructions (unless specified otherwise). 1 μg of total RNA was used for poly(A)-mRNA selection using streptavidin-coated magnetic beads. Then samples were fragmented to approximately 300bp and cDNA was synthesized using reverse transcriptase (SuperScript II, Invitrogen) and random primers. The second strand of the cDNA incorporated dUTP in place of dTTP. Double-stranded DNA was further used for library preparation. dsDNA was subjected to A-tailing and ligation of the barcoded Truseq adapters. All purification steps were performed using AMPure XP Beads (Agencourt). Library amplification was performed by PCR on the size selected fragments using the primer cocktail supplied in the kit. To estimate the quantity and check the fragment size libraries were analyzed using Agilent DNA 1000 chip (Agilent), and were subsequently quantified by qPCR using the KAPA Library Quantification Kit (KapaBiosystems) prior to amplification with Illumina’s cBot. Libraries were loaded and sequenced using 2×50 or 2×75 read lengths on Illumina’s HiSeq 2500. Experiments were performed in three biological replicates. All library preparation and sequencing steps were performed at the Genomics Unit of the Centre for Genomic Regulation (CRG), Barcelona, Spain.

### Bioinformatics analysis

### Quality control of sequencing data

We used FastQC v0.11.6 (http://www.bioinformatics.babraham.ac.uk/projects/fastqc) and Multiqc v. 1.0 (Ewels et al. 2016) to perform quality control of raw sequencing data.

### Phasing of heterozygous genomic regions

The reference genome and genome annotations for the putative parent strain *C. orthopsilosis* 90-125 were obtained from NCBI (assembly ASM31587v1, last accessed on 12/08/2018). To assess allele-specific expression in the hybrid strains we phased the hybrid genomes following the procedure illustrated in Fig. 1 and further described below.

**Fig. 1.**
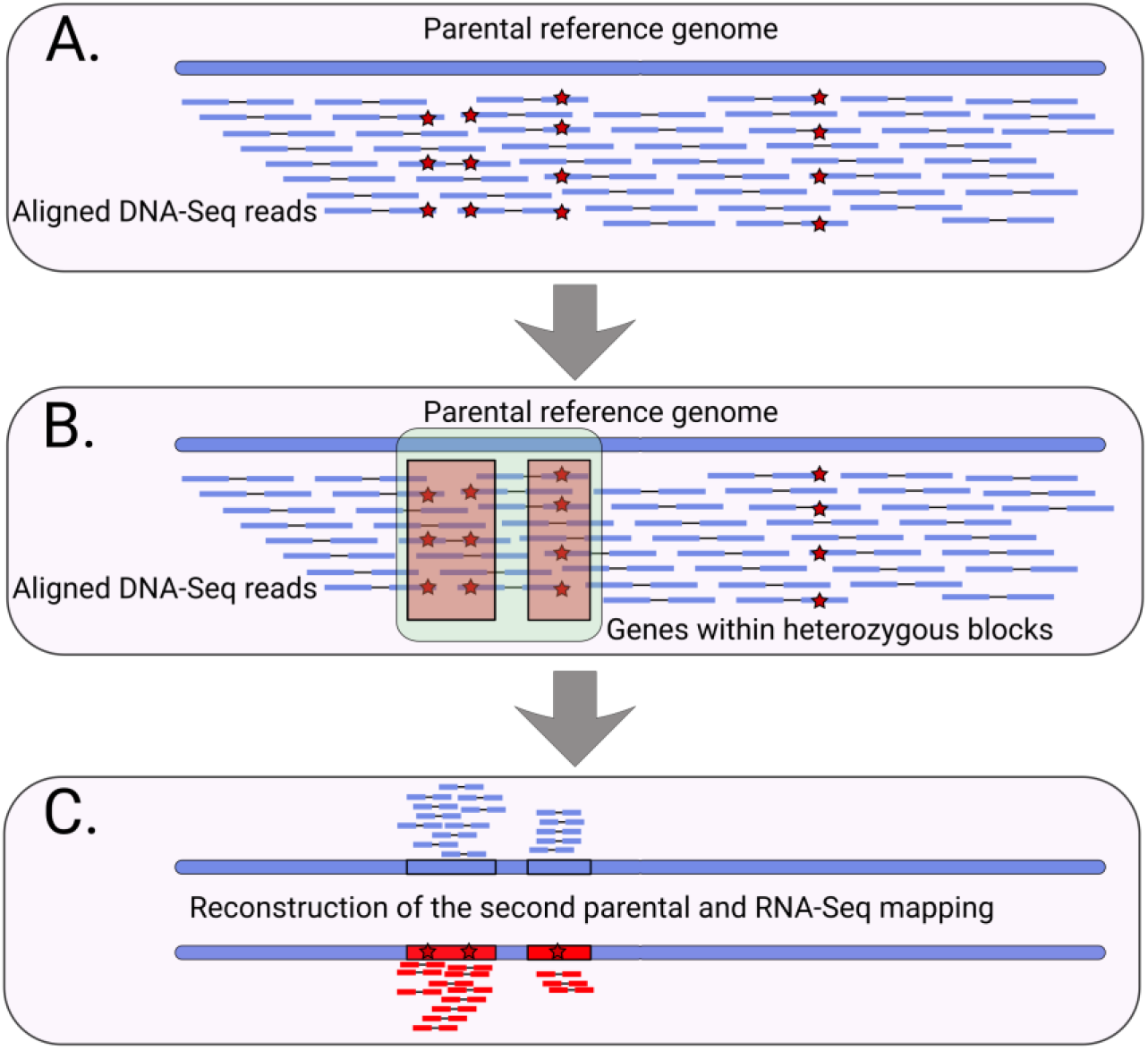
Schematic representation of the bioinformatics approach to assess the allele-specific expression in the hybrid strains. **(A)** Mapping of DNA-Seq reads to the parental reference genome and further variant calling (red stars represent heterozygous variants). **(B)** Defining heterozygous blocks (green rectangle) and identifying genes (red rectangles) within the blocks. **(C)** Inserting the heterozygous variants in the reference genome (second parental reconstruction) and further RNA-Seq read mapping to the partially phased genome.

Specifically, we first phased (i.e. resolved alternative haplotypes) the genes located in the heterozygous regions in the MCO456 and CP124 hybrid strains. To do this, we used DNA sequencing data of these strains (ERR321926 (Pryszcz et al. 2014, 2015) and SRR3547561 (Schröder et al. 2016)). We first trimmed these data using Trimmomatic v. 0.36 software with default parameters, and subsequently processed the data using mapping and variant calling modules of HaploTypo v.1 pipeline, setting the filter of SNP clusters at 5 SNPs in 20 bp window (Pegueroles et al. 2019), and subsequently removed indels using vcftools v0.1.16 (Danecek et al. 2011). Then using the heterozygous variants we defined heterozygous blocks using bedtools v2.29 (Quinlan and Hall 2010) with *merge* function as described in (Pryszcz et al. 2015) and further optimized in (Mixão et al. 2019) – if the distance between two heterozygous variants is less than 100 base pairs (bp), that region constitutes a heterozygous block, if the next variant to that block is located in less than 100 bp, the block is extended, otherwise it is interrupted. After defining heterozygous blocks we identified genes located within the blocks using a custom python script find_genes_in_heterozygous_blocks.py v1.0 (available at https://github.com/Gabaldonlab/C.-orthopsilosis-ASE/blob/master/find_genes_in_heterozygous_blocks.py). Samtools v. 1.9 was used to index the reference genome. Sorting of vcf files was done by *sort* function in bash. Subsequently, we inserted the alternative variants of the genes located in heterozygous blocks in the reference genome of the 90-125 strain using GATK v.3.7 (DePristo et al. 2011), thus reconstructing the sequence of the alternative parental genome within the defined heterozygous blocks.

### RNA-Seq and allele-specific expression analysis

RNA-Seq read mapping and summarization was performed using the splice-junction aware mapper STAR v. 2.7.3a (Dobin et al. 2012). GFF to GTF format conversion for genome annotations was done using gffread v. 0.11.6 (Trapnell et al. 2012) utility. For 90-125 strain, we mapped RNA-Seq data to the 90-125 reference assembly, while for the hybrid strains the data were mapped to a concatenated reference genomes, obtained by combining the 90-125 reference and the reconstructed parental reference. For mapping to the concatenated reference we set STAR option --outFilterMismatchNmax to 0 to restrict the mismatches in read alignments.

Differential gene expression and allele-specific expression were assessed using DESeq2 v. 1.22.2 (Love, Huber, and Anders 2014). For allele-specific expression comparisons the sizeFactors were set to 1 for all the samples, since the read counts for alleles come from the same library. For a gene to be considered differentially (allele-specifically) expressed, we used a threshold of |log2 fold change (L2FC)| > 1.5 and padj (adjusted p-value) < 0.01. Hypergeometric tests were performed using phyper function of R with lower.tail and log.p parameters set to FALSE.

To visualize gene expression data we used ggplot2 v. 2_3.0.0 R library (Wickham 2016). Putative functions of *C. orthopsilosis* genes were retrieved from *Candida* Genome Database (last accessed on 26 July 2019, Skrzypek et al. 2017).

RNA-Seq data is deposited at the SRA database under the accession numbers SRR10251160-SRR10251168.

## Results and Discussion

To understand the impact of hybridization on gene expression, and disentangle the transcriptomic interactions between the two parental subgenomes in the pathogenic hybrid yeast *C. orthopsilosis,* we performed RNA-Seq of two hybrid strains, and a homozygous strain belonging to one of the putative parental lineages (Pryszcz et al. 2014; Schröder et al. 2016). Importantly, the two analyzed hybrid strains belong to two independently formed hybrid clades resulting from the mating of the same two parental lineages: strain MCO456 belongs to the hybrid clade 1, which underwent extensive LOH, whereas strain CP124 belongs to clade 4, which has limited LOH (Fig. 2). Additionally, we analyzed a strain belonging to one of the putative parental homozygous lineages (90-125) and compared its transcriptomic profile with the hybrids. The summary statistics of our RNA-Seq data including mapping rates and reproducibility metrics are available in supplementary tables S1 and S2, respectively.

**Fig. 2.**
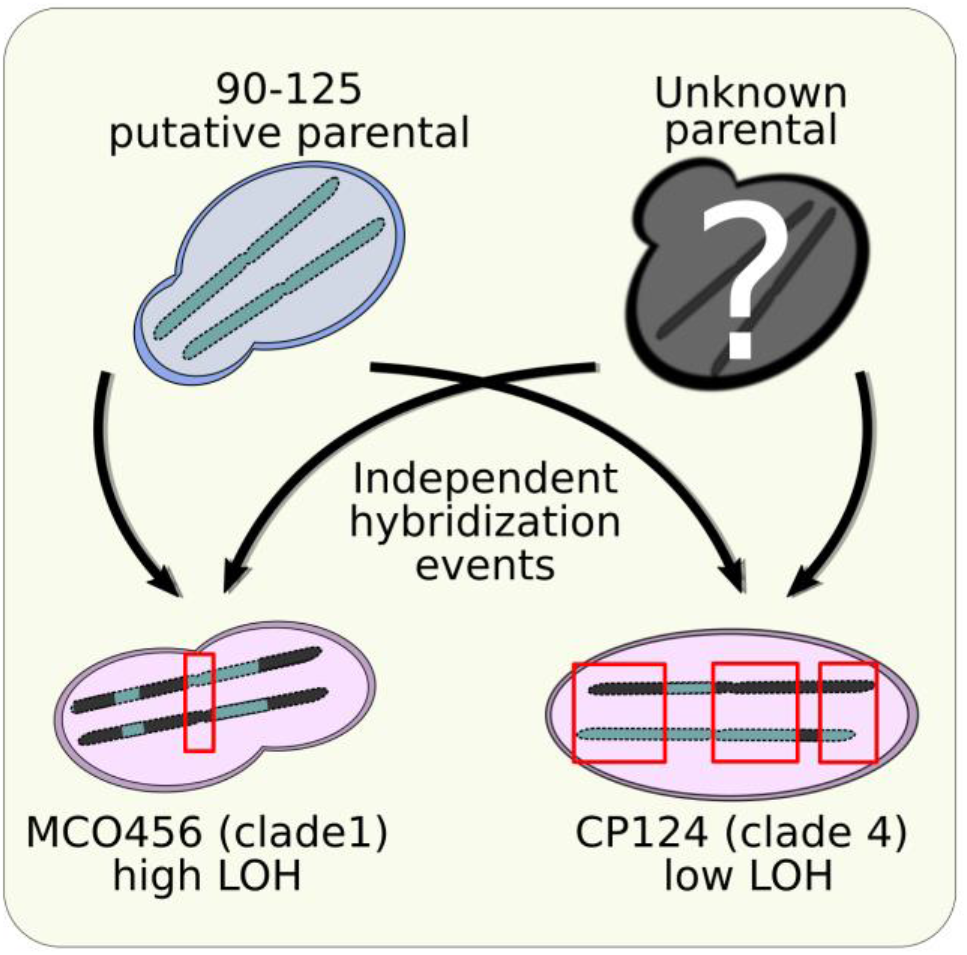
Schematic representation of the experimental design of the study. The strain *C. orthopsilosis* 90-125 represents a putative parental lineage, which has undergone several independent hybridization events (black arrows) by mating with a second, unknown parental. A supposedly more ancient hybridization event has given rise to a hybrid clade including the MCO456 strain (“high LOH”), which experienced extensive LOH, and a more recent hybridization event has lead to the formation of an independent hybrid clade including CP124 strain (“low LOH”), which contains more heterozygous regions (highlighted with red rectangles).

To perform accurate assignment of RNA-Seq reads to each of the two parental subgenomes in the hybrid, we used the following genome phasing procedure (see Figure 1 and Materials and Methods for details). Knowing the genome of a relative of one of the putative parentals (i.e. strain 90-125), we used it as a reference to map publicly available DNA-Seq raw data of the two hybrid strains. We phased the heterozygous regions in each of the hybrids by reconstructing the haplotype belonging to the known parental lineage (i.e. having the alleles present in the 90-125 sequence) and the alternative haplotype belonging to the unknown parental lineage (i.e having the heterozygous alleles alternative to 90-125). Using this procedure, we could phase 107 and 590 genes within heterozygous regions for MCO456 and CP124 strains, respectively (supplementary tables S3 and S4), from which 71 genes were common between the two strains. As expected, we obtained more phased genes in the clade 4 strain (CP124), which encompasses more heterozygous regions than MCO456 (Clade 1). Notably, the overlap of 71 genes in heterozygous regions between the two strains is more than expected by chance as calculated by hypergeometric test (p=1.011097e-45), which suggests the existence of structural or selective constraints acting on LOH events.

Once we obtained the phasing information, we further assessed the levels of allelic expression in hybrids by mapping the RNA-Seq data to the concatenated phased genomes using strict filters for read mapping mismatches (see Materials and Methods). We checked the rates of cross-mapping - i.e. reads that can not be unambiguously mapped to either parental - by mapping the data from the parental strain to the concatenated phased genomes, and observed a negligible proportion of cross-mapping in phased genes (~0.019% and ~0.023% for MCO456 and CP124, respectively, supplementary fig. S1 and S2). The expression levels in 90-125 strain were obtained by mapping RNA-Seq data directly to the reference genome. Read counts for all strains can be found in supplementary tables S5 and S6. Then we performed differential expression (DE) and allele-specific expression (ASE) analysis in both hybrid strains and the parental strain (Fig. 3).

**Fig. 3.**
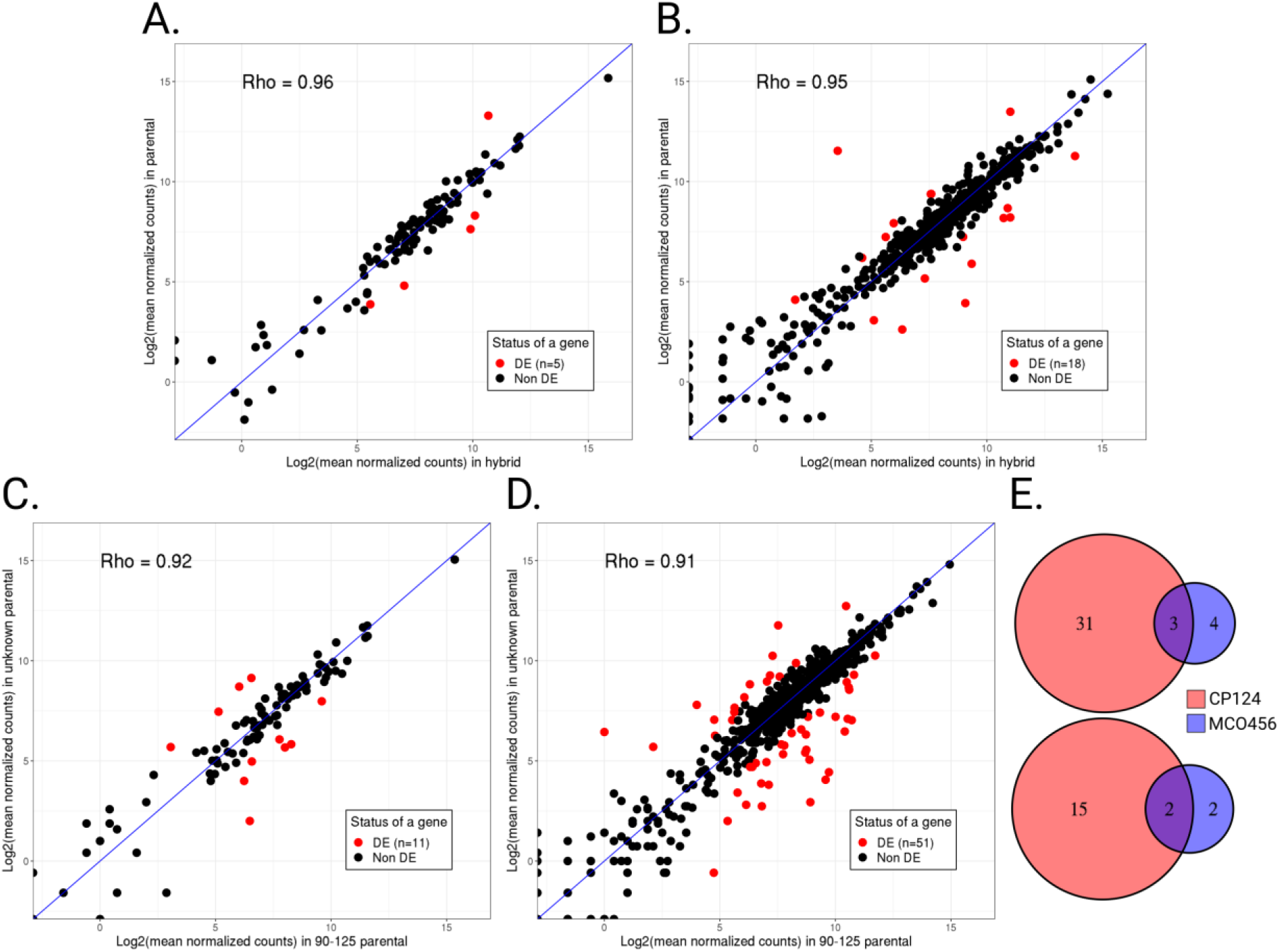
Overall results of DE and ASE comparisons. (**A)** Correlation between gene expression levels in 90-125 and MCO456 (**B)** Correlation between gene expression levels in 90-125 and CP124. **(C)** ASE analysis for *C. orthopsilosis* MCO456 strain **(D)** ASE analysis for CP124 strain. Scatter plots are based on mean normalized read counts for each gene. “DE” denotes **D**ifferentially **E**xpressed. **(E)** Venn diagrams showing the overlap between ASE genes in both strains: up-regulated 90-125 homeologs (on the top) and up-regulated homeologs of unknown parent (at the bottom)

We first compared the expression levels of 90-125 genes with the corresponding genes in heterozygous blocks in the hybrid strains. Such parental-hybrid comparisons (Fig. 3A and B) revealed a limited effect of hybridization on the parental gene expression levels - 5 (4.6%) and 18 (3%) DE genes when comparing 90-125 with MCO456 and CP124 hybrid, respectively (supplementary tables S7 and S8). Most of the differentially expressed genes have unknown roles, with some exceptions (supplementary tables S9 and S10). For example, CORT_0B00460 which is up-regulated in the hybrid MCO456 background, is predicted to have functions related to metal ion binding and superoxide metabolic processes.

Further, we performed allele-specific gene expression analysis in the hybrids. We detected 11 allele-specifically expressed (ASE) genes in MCO456 (10.2% of phased genes, supplementary table S11), while CP124 showed 51 ASE genes (8.6% of phased genes, supplementary table S12). Putative functions for ASE genes in MCO456 and CP124 strains can be found in supplementary tables S13 and S14, respectively. Although the function of most ASE genes was unknown, some have orthologs involved in virulence in other *Candida* pathogens. For both strains, genes related to superoxide dismutase activity (Broxton and Culotta 2016; Martchenko et al. 2004; C. X. Li et al. 2015) (CORT_0B00460 in MCO456, and CORT_0A12390 in CP124 strain) were up-regulated for the allele of the unknown parent (Broxton and Culotta 2016; Martchenko et al. 2004; C. X. Li et al. 2015). When comparing the 90-125 parental with the corresponding homeologs in the hybrids, the expression level of CORT_0A12390 was intact upon hybridization, while CORT_0B00460 was up-regulated in MCO456.

Moreover, we identified ASE genes related to zinc metabolism, which is one important player in host-pathogen interactions (Jung 2015; Wilson, Citiulo, and Hube 2012; Crawford and Wilson 2015). The gene CORT_0C02470, potentially involved in zinc transmembrane transport, was up-regulated in the hybrid towards the 90-125 parental allele in MCO456, while CORT_0E02010, with a putative zinc binding activity, is expressed at higher levels from the allele assigned to the unknown parental in CP124. For both genes, the expression levels were not altered upon hybridization.

Notably, five genes - CORT_0A04580, CORT_0A09590, CORT_0A03280, CORT_0D04190, and CORT_0E01400, are common ASE genes between the two hybrids, which is higher than expected by chance considering the shared fraction of heterozygous genes (Fig. 3E, p=9.433106e-06). While the functions of the first four genes are unknown, the orthologs of CORT_0E01400 gene, which expresses more the unknown parental allele, are involved in cellular response to drugs and extracellular region localization (according to CGD annotations).

## Discussion

Here, we investigated the transcriptomic interactions of divergent parental sub-genomes in *C. orthopsilosis* hybrids. To our knowledge this represents the first such study in a hybrid human opportunistic yeast pathogen. Our experimental design allowed us not only to compare expression of genes in hybrid genetic background to that in homozygous parental background, but also to assess the extent of convergent evolution in two independently formed hybrid lineages.

In agreement with previous studies of transcriptomic shock in fungal hybrids (Cox et al. 2014; Hovhannisyan et al. 2020), our results indicate that hybridization has a rather moderate effect on gene expression levels – on average ~4% of the studied genes changed the expression levels upon hybridization, as compared to the parental. This relatively low levels of transcriptomic alteration upon hybridization in yeast hybrids, contrasts with the larger levels reported in animals or plants, as noted earlier (Hovhannisyan et al. 2020), and reinforcing the idea that yeasts have a comparatively higher capacity than plants and animals to buffer the effects of the transcriptomic shock elicited by hybridization. As more transcriptomic studies on diverse organisms accumulate, it will be clarified how widespread these differences are and what molecular mechanisms may underlie this phenomenon.

When comparing the results of this study with those of the report assessing transcriptomic shock in artificial yeast hybrid (Hovhannisyan et al. 2020), we noted that the proportion of DE genes (calculated over the total number of genes in heterozygous regions) in natural hybrids is somewhat larger (~4% in this study), than in the case of the artificial *S. cerevisiae x S. uvarum* hybrids (~1.5%), despite the larger parental divergence and more recent nature of the hybridization. Additionally, we found that the two independently formed *C. orthopsilosis* hybrids, retained a shared subset of genes in heterozygosis, which is larger than expected by chance. Altogether these results suggest that structural constraints or functional selection may play a role in shaping LOH patterns in these hybrids, with certain genes, including some showing divergent expression patterns, being more likely to be retained in heterozygosis. Moreover, it has been previously reported that LOH events can be driven by selective pressures (Smukowski Heil et al. 2017). A plausible scenario is that shared genes in heterozygosis, and shared DE genes between the two independently formed hybrids are involved in traits beneficial for the hybrid, and are thus maintained through purifying selection. Nevertheless, comparisons from such limited number of studies must be taken with caution, and we hope that future studies will help to clarify such questions.

When assessing differences in expression between homeologous copies in the hybrid (ASE), we found that the fraction of genes with ASE (8.6% in CP124, and 10.2% in MCO456) was comparatively larger than the fraction of DE genes. In addition, these fractions of ASE genes are slightly higher than that noted for a newly formed *S. cerevisiae x S. uvarum* hybrid (7.4%), despite the much lower parental divergence in *C. orthopsilosis* hybrids. Moreover, the fraction of ASE genes is larger in MCO456 (10.2% as compared to 8.6% in CP124) which underwent more extensive LOH. Although more datasets are needed to confirm this trend, this observation suggests that LOH preferentially targets genes that do not display ASE, consistent with the observation above for DE genes. Of note, we did not observe a strong preferential expression of alleles from either of parental subgenomes with 34 (~66%) and 7 (~63%) of ASE genes were expressed higher in the known parental in CP124 and MCO456 strains, respectively. This is in line with initial comparisons of *C. orthopsilosis* genomes, which showed no preferential retention of any of the parental genomes in regions undergoing LOH (Pryszcz et al. 2014). Previous studies in natural (*Epichloë*) and artificial (*S. cerevisiae x S. uvarum*) fungal hybrids have also reported no strong preferential over-expression of one of the parental sub-genomes (Cox et al. 2014; X. C. Li and Fay 2017), which suggests this may be a general phenomenon in ascomycete fungal hybrids.

For both *C. orthopsilosis* hybrid strains we identified ASE genes involved in processes directly related to virulence, such as zinc ion transport and superoxide dismutase activity. However, since the second parental organism is still unidentified it is unknown whether the observed expression divergence between homeologs arose due to hybridization or whether it was already existing between orthologs of the parental species. Although we do not claim that these transcriptomic differences are actually related to differences in virulence, we consider that this possibility should be considered in future studies.

## Conclusions

Altogether, our analyses provide evidence of a moderate effect of hybridization on the transcriptome of pathogenic hybrid yeasts, in line with previous observations in other fungal hybrids. Interestingly, the significant overlap of genes in heterozygous blocks, including DE and ASE genes, in the two independently formed hybrids suggests the existence of selecting constraints acting on genes that show altered expression in the hybrid. Similarly, we detected that increase in LOH was associated with higher fractions of ASE genes, suggesting that ASE genes are preferentially retained in heterozygosis in the hybrid. Finally, we detected ASE genes in the studied pathogen which are known to have direct implications in fungal virulence in other yeast species, like *C. albicans*, making them a target for further studies of fungal pathogenicity emergence.

## Acknowledgements

We would like to thank Verónica Mixão (BSC/IRB, Barcelona, Spain) for her help in performing analysis with the HaploTypo pipeline. This work received funding from the European Union’s Horizon 2020 Research and Innovation Programme under the Marie Skłodowska-Curie Grant Agreement No. H2020-MSCA-ITN-2014-642095. TG group also acknowledges support from the Spanish Ministry of Economy, Industry, and Competitiveness (MEIC) for the EMBL partnership, and grants ‘Centro de Excelencia Severo Ochoa 2013– 2017’ SEV-2012-0208, and BFU2015-67107 co-founded by European Regional Development Fund (ERDF); from the CERCA Programme/Generalitat de Catalunya; from the Catalan Research Agency (AGAUR) SGR857, and grants from the European Union’s Horizon 2020 Research and Innovation Programme under the Grant Agreement No. ERC-2016-724173. TG also receives support from a INB grant (PT17/0009/0023 – ISCIII-SGEFI/ERDF).

## References

Boekhout, T., B. Theelen, M. Diaz, J. W. Fell, W. C. Hop, E. C. Abeln, F. Dromer, and W. Meyer. 2001. “Hybrid Genotypes in the Pathogenic Yeast Cryptococcus Neoformans.” Microbiology 147 (Pt 4): 891–907.

Bovers, Marjan, Ferry Hagen, Eiko E. Kuramae, Hans L. Hoogveld, Françoise Dromer, Guy St-Germain, and Teun Boekhout. 2008. “AIDS Patient Death Caused by Novel Cryptococcus Neoformans X C. Gattii Hybrid.” Emerging Infectious Diseases 14 (7): 1105–8.

Broxton, Chynna N., and Valeria C. Culotta. 2016. “SOD Enzymes and Microbial Pathogens: Surviving the Oxidative Storm of Infection.” PLoS Pathogens 12 (1): e1005295.

Callaghan, Sophia, and David Guest. 2015. “Globalisation, the Founder Effect, Hybrid Phytophthora Species and Rapid Evolution: New Headaches for Biosecurity.” Australasian Plant Pathology. https://doi.org/10.1007/s13313-015-0348-5.

Cox, Murray P., Ting Dong, Genggeng Shen, Yogesh Dalvi, D. Barry Scott, and Austen R. D. Ganley. 2014. “An Interspecific Fungal Hybrid Reveals Cross-Kingdom Rules for Allopolyploid Gene Expression Patterns.” PLoS Genetics 10 (3): e1004180.

Crawford, Aaron, and Duncan Wilson. 2015. “Essential Metals at the Host-Pathogen Interface: Nutritional Immunity and Micronutrient Assimilation by Human Fungal Pathogens.” FEMS Yeast Research 15 (7). https://doi.org/10.1093/femsyr/fov071.

Danecek, Petr, Adam Auton, Goncalo Abecasis, Cornelis A. Albers, Eric Banks, Mark A. DePristo, Robert E. Handsaker, et al. 2011. “The Variant Call Format and VCFtools.” Bioinformatics 27 (15): 2156–58.

Depotter, Jasper R. L., Silke Deketelaere, Patrik Inderbitzin, Andreas Von Tiedemann, Monica Höfte, Krishna V. Subbarao, Thomas A. Wood, and Bart P. H. J. Thomma. 2016. “Verticillium Longisporum, the Invisible Threat to Oilseed Rape and Other Brassicaceous Plant Hosts.” Molecular Plant Pathology 17 (7): 1004–16.

Depotter, Jasper Rl, Michael F. Seidl, Thomas A. Wood, and Bart Phj Thomma. 2016. “Interspecific Hybridization Impacts Host Range and Pathogenicity of Filamentous Microbes.” Current Opinion in Microbiology 32 (August): 7–13.

DePristo, Mark A., Eric Banks, Ryan Poplin, Kiran V. Garimella, Jared R. Maguire, Christopher Hartl, Anthony A. Philippakis, et al. 2011. “A Framework for Variation Discovery and Genotyping Using next-Generation DNA Sequencing Data.” Nature Genetics 43 (5): 491–98.

Dobin, Alexander, Carrie A. Davis, Felix Schlesinger, Jorg Drenkow, Chris Zaleski, Sonali Jha, Philippe Batut, Mark Chaisson, and Thomas R. Gingeras. 2012. “STAR: Ultrafast Universal RNA-Seq Aligner.” Bioinformatics 29 (1): 15–21.

Dutta, Abhishek, Gen Lin, Ajith V. Pankajam, Parijat Chakraborty, Nahush Bhat, Lars M. Steinmetz, and Koodali T. Nishant. 2017. “Genome Dynamics of Hybrid Saccharomyces Cerevisiae During Vegetative and Meiotic Divisions.” G3: Genes|Genomes|Genetics. https://doi.org/10.1534/g3.117.1135.

Ewels, Philip, Måns Magnusson, Sverker Lundin, and Max Käller. 2016. “MultiQC: Summarize Analysis Results for Multiple Tools and Samples in a Single Report.” Bioinformatics 32 (19): 3047–48.

Gabaldón, Toni, Miguel A. Naranjo-Ortíz, and Marina Marcet-Houben. 2016. “Evolutionary Genomics of Yeast Pathogens in the Saccharomycotina.” FEMS Yeast Research 16 (6). https://doi.org/10.1093/femsyr/fow064.

Garcia-Solache, Monica A., and Arturo Casadevall. 2010. “Global Warming Will Bring New Fungal Diseases for Mammals.” mBio 1 (1). https://doi.org/10.1128/mBio.00061-10.

Greaves, Ian K., Rebeca Gonzalez-Bayon, Li Wang, Anyu Zhu, Pei-Chuan Liu, Michael Groszmann, W. James Peacock, and Elizabeth S. Dennis. 2015. “Epigenetic Changes in Hybrids.” Plant Physiology. https://doi.org/10.1104/pp.15.00231.

Hovhannisyan, Hrant, Ester Saus, Ewa Ksiezopolska, Alex J Hinks Roberts, Edward J. Louis, Toni Gabaldón. 2020. “Integrative Omics Analysis Reveals a Limited Transcriptional Shock After Yeast Inter-species Hybridization.” Frontiers in Genetics (in press).

Hu, Xiaojiao, Hongwu Wang, Kun Li, Yujin Wu, Zhifang Liu, and Changling Huang. 2017. “Genome-Wide Proteomic Profiling Reveals the Role of Dominance Protein Expression in Heterosis in Immature Maize Ears.” Scientific Reports 7 (1): 16130.

Jonge, R. de, R. de Jonge, M. D. Bolton, A. Kombrink, G. C. M. van den Berg, K. A. Yadeta, and B P H. 2013. “Extensive Chromosomal Reshuffling Drives Evolution of Virulence in an Asexual Pathogen.” Genome Research. https://doi.org/10.1101/gr.152660.112.

Jung, Won Hee. 2015. “The Zinc Transport Systems and Their Regulation in Pathogenic Fungi.” Mycobiology 43 (3): 179–83.

Ksiezopolska, Ewa, and Toni Gabaldón. 2018. “Evolutionary Emergence of Drug Resistance in Candida Opportunistic Pathogens.” Genes 9 (9). https://doi.org/10.3390/genes9090461.

Li, Cissy X., Julie E. Gleason, Sean X. Zhang, Vincent M. Bruno, Brendan P. Cormack, and Valeria Cizewski Culotta. 2015. “Candida Albicans Adapts to Host Copper during Infection by Swapping Metal Cofactors for Superoxide Dismutase.” Proceedings of the National Academy of Sciences of the United States of America 112 (38): E5336–42.

Li, Xueying C., and Justin C. Fay. 2017. “Cis-Regulatory Divergence in Gene Expression between Two Thermally Divergent Yeast Species.” Genome Biology and Evolution. https://doi.org/10.1093/gbe/evx072.

Love, Michael I., Wolfgang Huber, and Simon Anders. 2014. “Moderated Estimation of Fold Change and Dispersion for RNA-Seq Data with DESeq2.” Genome Biology 15 (12): 550.

Marcet-Houben, Marina, and Toni Gabaldón. 2015. “Beyond the Whole-Genome Duplication: Phylogenetic Evidence for an Ancient Interspecies Hybridization in the Baker’s Yeast Lineage.” PLOS Biology. https://doi.org/10.1371/journal.pbio.1002220.

Martchenko, Mikhail, Anne-Marie Alarco, Doreen Harcus, and Malcolm Whiteway. 2004. “Superoxide Dismutases in Candida Albicans: Transcriptional Regulation and Functional Characterization of the Hyphal-Induced SOD5 Gene.” Molecular Biology of the Cell 15 (2): 456–67.

McClintock, B. 1984. “The Significance of Responses of the Genome to Challenge.” Science. https://doi.org/10.1126/science.15739260.

Metzger, Brian P. H., Patricia J. Wittkopp, and Joseph D. Coolon. 2017. “Evolutionary Dynamics of Regulatory Changes Underlying Gene Expression Divergence among Saccharomyces Species.” Genome Biology and Evolution 9 (4): 843–54.

Mixão, Verónica, and Toni Gabaldón. 2018. “Hybridization and Emergence of Virulence in Opportunistic Human Yeast Pathogens.” Yeast 35 (1): 5–20.

Mixão, Verónica, Antonio Perez Hansen, Ester Saus, Teun Boekhout, Cornelia Lass-Florl, and Toni Gabaldón. 2019. “Whole-Genome Sequencing of the Opportunistic Yeast Pathogen Candida Inconspicua Uncovers Its Hybrid Origin.” Frontiers in Genetics. https://doi.org/10.3389/fgene.2019.00383.

Pammi, Mohan, Linda Holland, Geraldine Butler, Attila Gacser, and Joseph M. Bliss. 2013. “Candida Parapsilosis Is a Significant Neonatal Pathogen: A Systematic Review and Meta-Analysis.” The Pediatric Infectious Disease Journal 32 (5): e206–16.

Papon, Nicolas, Vincent Courdavault, Marc Clastre, and Richard J. Bennett. 2013. “Emerging and Emerged Pathogenic Candida Species: Beyond the Candida Albicans Paradigm.” PLoS Pathogens 9 (9): e1003550.

Pegueroles, Cinta, Verónica Mixão, Laia Carreté, Manu Molina, and Toni Gabaldón. 2019. “HaploTypo: A Variant-Calling Pipeline for Phased Genomes.” Bioinformatics, December. https://doi.org/10.1093/bioinformatics/btz933.

Pfaller, M. A., and D. J. Diekema. 2007. “Epidemiology of Invasive Candidiasis: A Persistent Public Health Problem.” Clinical Microbiology Reviews 20 (1): 133–63.

Pfaller, M. A., S. A. Messer, R. J. Hollis, L. Boyken, S. Tendolkar, J. Kroeger, and D. J. Diekema. 2009. “Variation in Susceptibility of Bloodstream Isolates of Candida Glabrata to Fluconazole according to Patient Age and Geographic Location in the United States in 2001 to 2007.” Journal of Clinical Microbiology 47 (10): 3185–90.

Pryszcz, Leszek P., Tibor Németh, Attila Gácser, and Toni Gabaldón. 2014. “Genome Comparison of Candida Orthopsilosis Clinical Strains Reveals the Existence of Hybrids between Two Distinct Subspecies.” Genome Biology and Evolution 6 (5): 1069–78.

Pryszcz, Leszek P., Tibor Németh, Ester Saus, Ewa Ksiezopolska, Eva Hegedűsová, Jozef Nosek, Kenneth H. Wolfe, Attila Gacser, and Toni Gabaldón. 2015. “The Genomic Aftermath of Hybridization in the Opportunistic Pathogen Candida Metapsilosis.” PLoS Genetics 11 (10): e1005626.

Quinlan, Aaron R., and Ira M. Hall. 2010. “BEDTools: A Flexible Suite of Utilities for Comparing Genomic Features.” Bioinformatics 26 (6): 841–42.

Riccombeni, Alessandro, Genevieve Vidanes, Estelle Proux-Wéra, Kenneth H. Wolfe, and Geraldine Butler. 2012. “Sequence and Analysis of the Genome of the Pathogenic Yeast Candida Orthopsilosis.” PloS One 7 (4): e35750.

Schröder, Markus S., Kontxi Martinez de San Vicente, Tâmara H. R. Prandini, Stephen Hammel, Desmond G. Higgins, Eduardo Bagagli, Kenneth H. Wolfe, and Geraldine Butler. 2016. “Multiple Origins of the Pathogenic Yeast Candida Orthopsilosis by Separate Hybridizations between Two Parental Species.” PLoS Genetics 12 (11): e1006404.

Skrzypek, Marek S., Jonathan Binkley, Gail Binkley, Stuart R. Miyasato, Matt Simison, and Gavin Sherlock. 2017. “The Candida Genome Database (CGD): Incorporation of Assembly 22, Systematic Identifiers and Visualization of High Throughput Sequencing Data.” Nucleic Acids Research 45 (D1): D592–96.

Smukowski Heil, Caiti S., Christopher G. DeSevo, Dave A. Pai, Cheryl M. Tucker, Margaret L. Hoang, and Maitreya J. Dunham. 2017. “Loss of Heterozygosity Drives Adaptation in Hybrid Yeast.” Molecular Biology and Evolution 34 (7): 1596–1612.

Tavanti, Arianna, Amanda D. Davidson, Neil A. R. Gow, Martin C. J. Maiden, and Frank C. Odds. 2005. “Candida Orthopsilosis and Candida Metapsilosis Spp. Nov. to Replace Candida Parapsilosis Groups II and III.” Journal of Clinical Microbiology 43 (1): 284–92.

Tirosh, Itay, Sharon Reikhav, Avraham A. Levy, and Naama Barkai. 2009. “A Yeast Hybrid Provides Insight into the Evolution of Gene Expression Regulation.” Science 324 (5927): 659–62.

Tóth, Renáta, Jozef Nosek, Héctor M. Mora-Montes, Toni Gabaldon, Joseph M. Bliss, Joshua D. Nosanchuk, Siobhán A. Turner, Geraldine Butler, Csaba Vágvölgyi, and Attila Gácser. 2019. “Candida Parapsilosis: From Genes to the Bedside.” Clinical Microbiology Reviews 32 (2). https://doi.org/10.1128/CMR.00111-18.

Trapnell, Cole, Adam Roberts, Loyal Goff, Geo Pertea, Daehwan Kim, David R. Kelley, Harold Pimentel, Steven L. Salzberg, John L. Rinn, and Lior Pachter. 2012. “Differential Gene and Transcript Expression Analysis of RNA-Seq Experiments with TopHat and Cufflinks.” Nature Protocols 7 (3): 562–78.

Wickham, Hadley. 2016. “Programming with ggplot2.” In Use R!, 241–53.

Wilson, Duncan, Francesco Citiulo, and Bernhard Hube. 2012. “Zinc Exploitation by Pathogenic Fungi.” PLoS Pathogens 8 (12): e1003034.

Wu, Guangxi, He Zhao, Chenhao Li, Menaka Priyadarsani Rajapakse, Wing Cheong Wong, Jun Xu, Charles W. Saunders, et al. 2015. “Genus-Wide Comparative Genomics of Malassezia Delineates Its Phylogeny, Physiology, and Niche Adaptation on Human Skin.” PLoS Genetics 11 (11): e1005614.

